# Madagascar as an evolutionary source for the Paleotropical flora: New insights from the biogeography of *Grewia* (Malvaceae)

**DOI:** 10.64898/2026.09.02.748941

**Authors:** Lucile Jourdain-Fievet, Margaret M. Hanes, Nisa Karimi, Laurence J. Dorr, Kenneth J. Wurdack

## Abstract

Oceanic islands are natural laboratories for investigating the processes that generate, maintain, and redistribute biodiversity through time. An ideal flowering plant system to investigate these processes is *Grewia* L. (Malvaceae, Grewioideae), comprising ca. 283 species, of which nearly 30% are endemic to Madagascar, where they display considerable morphological diversity. Using Angiosperms353 target-capture sequencing for 171 accessions (119 *Grewia* spp.), we reconstructed the first densely sampled phylogeny of *Grewia* and performed divergence-time estimation, diversification analyses, and ancestral-range reconstruction. Our results support a Malagasy origin of the crown group around 27 million years ago, followed by extensive in situ diversification of *Grewia*. Repeated dispersal events from Madagascar to continental Africa subsequently contributed to the establishment of new African lineages, whereas colonization of the western Indian Ocean islands was more recent and resulted in anagenetic dispersals. Africa acted as the principal source of dispersal toward Arabia and into tropical Asia, suggesting the presence of a Miocene Afro-Arabian migration corridor under warm and humid conditions. Subsequent eastward dispersal from Asia to Australia is consistent with the stepping stone pathway provided by the Sunda and Sahul regions during the Miocene. This complex dispersal history was likely aided by fleshy fruits adapted for endozoochory. Our new evolutionary framework provides a robust basis for future taxonomic revision of this morphologically and ecologically diverse genus, and for identifying which characters have contributed to its exceptional diversification.

**Highlights:**

This is the first phylogenomic analysis of *Grewia*, one of the largest genera in Malvaceae.
We reconstruct dispersal events for *Grewia* across the Paleotropics.
Miocene corridors and fleshy fruits aided dispersal out of Africa and eastward.
*Grewia* originated in Madagascar at 27 Ma and colonised Africa at least 5 times.
Madagascar is the center of *Grewia* diversity with more than 80 endemic species.

## 1. Introduction

A central question in historical biogeography is understanding the origin and evolution of biodiversity hotspots. Oceanic islands, with their remarkable levels of species richness and endemism are often such hotspots and have long fascinated biologists as natural laboratories for studying the processes that generate, maintain, and redistribute biodiversity through time (MacArthur and Wilson, 1967; Whittaker et al., 2008, 2017; Gillespie and Roderick, 2014; Matthews and Triantis, 2021). Present-day diversity patterns result from the interplay of dispersal, vicariance, in situ diversification, extinction, climatic fluctuations, and geological processes operating over millions of years. Reconstructing the relative importance of these mechanisms has become a major objective of modern biogeography (Crisp et al., 2011).

Madagascar occupies a unique position among tropical biodiversity hotspots. It harbours one of the richest and most endemic floras worldwide (Myers et al., 2000; Antonelli et al., 2022) despite representing less than 0.4% of the Earth’s terrestrial surface. Current estimates indicate that the island supports approximately 14,900 native vascular plant species, of which more than 87% are endemic (Callmander et al., 2011; Antonelli et al., 2022). This remarkable diversity has traditionally been attributed to Madagascar’s long geological isolation and pronounced environmental heterogeneity over small geographic distances that together have promoted extensive in situ diversification across numerous plant and animal lineages (Yoder and Nowak, 2006; Goodman, 2022).

For decades, the origin of Madagascar’s biota was largely interpreted through the lens of Gondwanan vicariance following supercontinent fragmentation (Raven, 1979; Schatz, 1996). However, advances in phylogenetics and molecular dating have significantly altered this interpretation. A growing body of evidence from family Malvaceae now indicates that many Malagasy lineages are younger than the geological separation of Madagascar from India (Richardson et al., 2015; Hernández-Gutiérrez and Magallón, 2019; Cvetković et al., 2021), implying repeated oversea dispersals followed by diversification within the island (Koopman and Baum, 2008; Le Péchon et al., 2010; Skema and Hanes, 2022; Skema et al., 2023). Although the timing of colonization by plants on Madagascar has received some attention, the relative importance of Madagascar as a source of widespread paleotropical diversity remains less well understood (though see Skema et al., 2023). Furthermore, quantifying the respective contributions of vicariance, long-distance dispersal, range expansion, and in situ diversification in widespread clades throughout the Paleotropics requires densely sampled, time-calibrated phylogenies spanning the entire distribution of a group.

An excellent flowering plant system in which to investigate these questions is *Grewia* L. (Malvaceae, Grewioideae). The genus comprises ca. 283 currently accepted (plus many undescribed) species (WFO, 2026) of trees, shrubs, or climbers with flowers bearing numerous stamens, 2–12 ovules, and drupaceous fruit that can be fleshy and edible (Fig. 1). *Grewia* spp. display wide morphological variation in their leaves (e.g., shape, size, trichomes) and reproductive features (i.e., flower color, stamen number, ovule number, fruit fleshiness). Species are found across a diversity of habitats from extreme arid scrublands to tropical rainforests. The genus is widely distributed throughout the Paleotropics (Fig. 2), extending from Madagascar and sub-Saharan Africa to the Arabian Peninsula, tropical Asia, Australia, New Guinea, and several Pacific islands. Two thirds of *Grewia* spp. occur in Madagascar and Africa. Madagascar is particularly diverse and has been proposed as the center of *Grewia* diversity, with 93 recorded species (of which 80, representing nearly 30% of the entire genus, are endemic) (Capuron, 1964; Capuron and Mabberley, 1999; Mabberley, 1999; Karimi and Hanes, 2023).

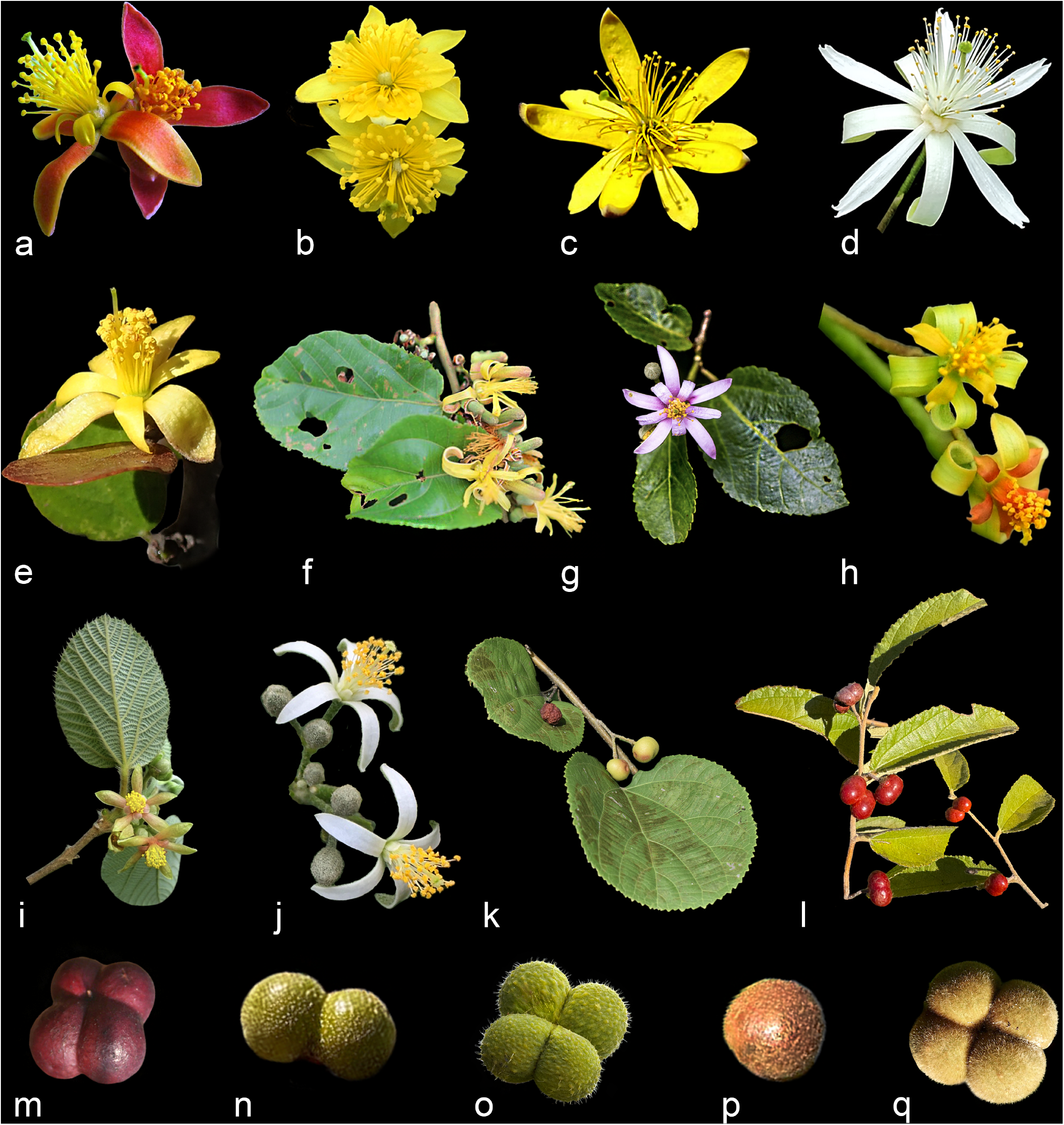

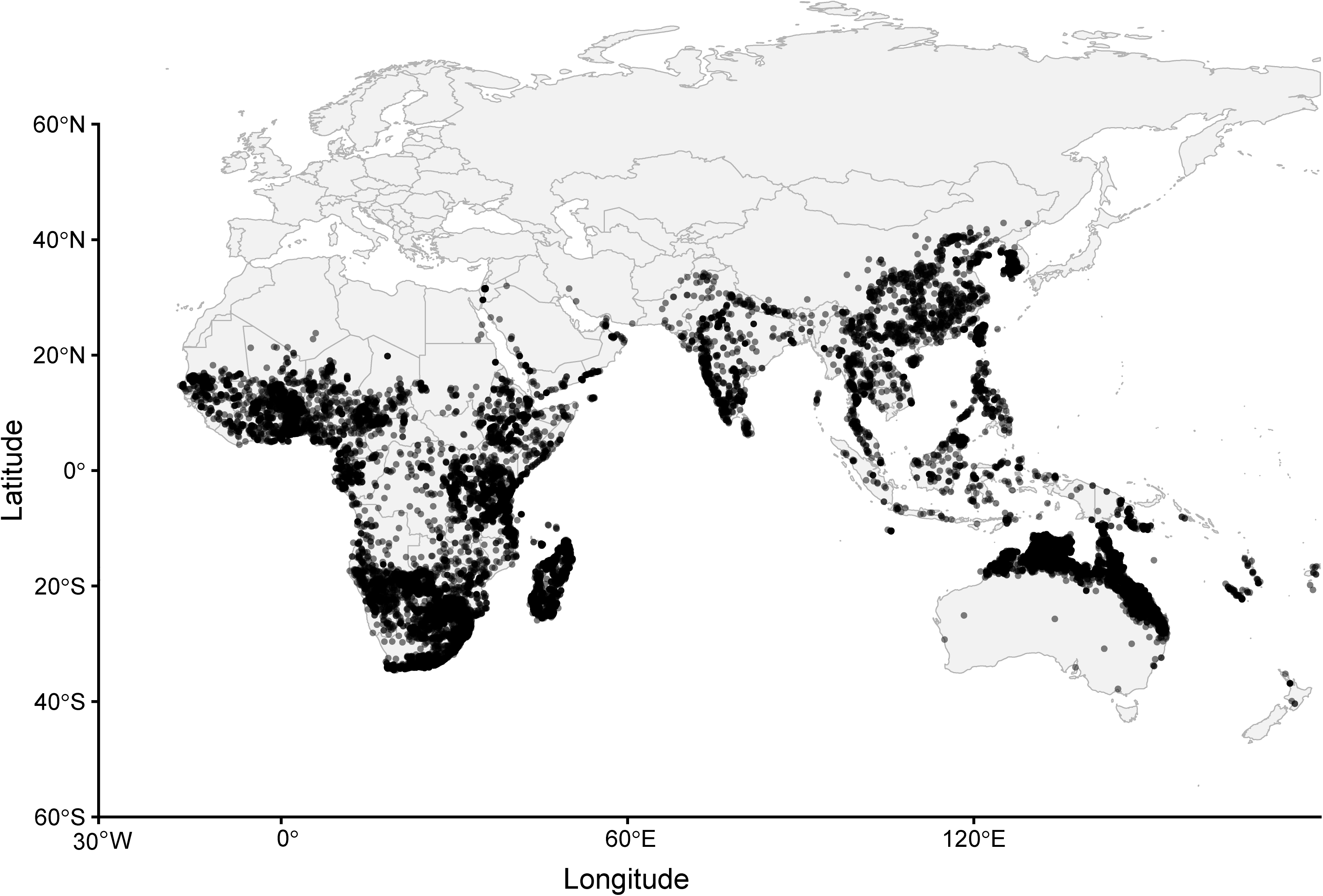

The absence of a comprehensive taxonomic revision of *Grewia* has long hindered a stable infrageneric classification for the genus. Species delimitation is challenging because of high species richness, extreme morphological variation and poorly defined species boundaries, particularly among the Malagasy lineages. Nevertheless, new species continue to be described based on morphological evidence (Wahlert et al. 2014, 2015; Mabberly and Capuron, 2017; Barrett, 2019; Lin et al., 2025; Phengmala et al., 2025; Ramon and Phillipson, 2026). The evolutionary history of *Grewia* also remains poorly understood. Prior molecular phylogenetic studies of relevance to resolving subfamily Grewioideae have sampled few species of *Grewia* and/or few genetic loci (e.g., Brunken and Muellner, 2012; Richardson et al., 2015; Hoorn et al., 2019; Dorr and Wurdack, 2024; Colli-Silva et al., 2025; Yang et al., 2025). Genomic resources for the genus are sparse and include a handful of chloroplast genomes (Xu et al., 2021; Hou et al., 2023; Al-Juhani 2025) and a transcriptome (Jain et al., 2025), but no whole genome assembly.

The striking concentration of endemic diversity in Madagascar, combined with broad continental distributions of the genus, makes *Grewia* particularly well suited for investigating the historical assembly of the Paleotropical flora. Here, we present the first densely sampled phylogenomic reconstruction of *Grewia* across its geographic distribution using Angiosperms353 capture-enrichment baits. By integrating phylogenomics, divergence-time estimation, diversification analyses, ancestral-area reconstruction, and biogeographic stochastic mapping, we investigate the historical processes responsible for the diversification of one of the largest genera in Malvaceae. Specifically, we address three questions: (1) Where did *Grewia* originate? (2) When did its principal lineages diversify? and (3) What roles have dispersal, range expansion, vicariance, and in situ diversification played in shaping its present-day distribution?

## 2. Methods

### 2.1. Taxon sampling and laboratory methods

In this study, we analysed 171 accessions representing 143 taxa of Grewioideae, including 119 species of *Grewia* (42% of the genus) of which 106 are currently accepted species and 13 provisionally named Malagasy species (i.e., the names are in use but not yet validly published). Grewioideae presently contains 24 genera divided among two tribes, and our sampling, while generically incomplete, was strategic to test the monophyly of *Grewia*. The ultimate outgroup, *Mortoniodendron* Standl. & Steyerm., was placed in Tilioideae by Colli-Silva et al. (2025). Voucher information and provenance data for all accessions are provided in Supplementary File 1.

Total genomic DNAs were extracted from dried leaf fragments using a modified CTAB method (Doyle and Doyle, 1987; Johnson et al., 2023). High molecular weight DNA extracts (usually from silica-gel dried leaves) were sheared to a 300–500 bp length using a Q800R3 Sonicator (Qsonica, Newtown, CT, USA); degraded, herbarium specimen derived extracts omitted this shearing step. Samples with a low DNA concentration (< 40 ng / µL), were concentrated using an Eppendorf Vacufuge Plus (Hamburg, Germany) at 45° C for ca. 2 h (V-AQ option). Libraries were prepared with the KAPA HyperPrep Kit (Hoffmann-La Roche, Basel, Switzerland) following the manufacturer’s protocol with 500–1000 ng gDNA with custom-synthesized adaptors and index primers (Glenn et al., 2019) for a ¼ reaction of 13 µL per sample. Twelve indexed, normalized (60–300 ng of DNA per sample) libraries were pooled per hybrid enrichment reaction that was then hybridized with the Angiosperms353 probe set (Johnson et al., 2019) for 40 h at 65° C, following the myBaits v5.03 standard protocol (Arbor Biosciences, Ann Arbor, MI, USA). These enriched libraries were amplified using KAPA HiFi HotStart Ready Mix (Hoffmann-La Roche, Basel, Switzerland) with 19 cycles and purified with 0.9x KAPA Pure Beads. Finally, all amplified enrichments were combined at equal proportions in one pool and sequenced by Novogene (Sacramento, CA, USA) as 2 x 150 bp, paired-end reads on one lane of an Illumina Nova-Seq X Plus 10B.

### 2.3. Sequence data processing

We filtered low–quality reads and deleted adaptors using Trimmomatic v.0.39 (Bolger et al., 2014). TruSeq3-PE adapters were removed allowing a maximum of 2 bp mismatches, with a simple palindrome threshold of 30 bp, and a simple clip threshold of 10 bp. The read ends were trimmed of three bases when the Phred quality score was < 33. We performed a 4-base sliding window to trim after the average quality fell below 15 and the read quality after trimming was checked using Fastp v.1.3.3. (Chen et al., 2018).

We mapped those trimmed reads to the 353 genes Angiosperm bait set target file using the HybPiper pipeline v.2.1.6 (Johnson et al., 2016) and all potential paralogs detected by HybPiper were discarded. Multiple sequence alignments (MSAs) were performed for each locus separately using MAFFT v.7.525 (Katoh and Standley, 2013) and trimmed using trimAl v.1.4 (Capella-Gutiérrez et al., 2009) with a gap threshold of 0.75 to remove ambiguously aligned positions.

### 2.4. Phylogenomic analyses

Maximum likelihood analyses were conducted using IQ-TREE v.2.3.1 (Minh et al., 2020a) on a concatenated alignment of 353 loci. Branch support was assessed using the SH-like approximate likelihood ratio test (SH-aLRT; Anisimova and Gascuel, 2006) with 1,000 replicates and ultrafast bootstrap approximation (UFBoot; Hoang et al., 2018) with 1,000 replicates. Maximum likelihood gene trees were inferred independently for each locus using IQ-TREE and subsequent species-tree inference was with ASTRAL-III v.5.7.8 under the multispecies coalescent model (Zhang et al., 2018). To further assess phylogenomic concordance, gene concordance factors (gCF) and site concordance factors (sCF) were calculated in IQ-TREE (Minh et al., 2020b). Gene concordance factors were estimated from the individual gene trees, whereas site concordance factors were estimated from the sequence alignments using 100 randomly sampled quartets per branch. Node support on the species tree was evaluated using local posterior probabilities (LPP; Sayyari and Mirarab, 2016) and normalized quartet scores, which represent the proportion of gene-tree quartets supporting the inferred relationship relative to the two alternative topologies. All support values are provided in Supplementary File 2.

### 2.5. Divergence time estimation

Four calibration constraints were applied within the subfamily Grewioideae (Table 1). The crown age of Grewioideae was constrained between 55 and 75 Million years ago (Ma) following Hernández-Gutiérrez and Magallón (2019), who in a Malvales-wide analysis inferred a crown age of 64.41 Ma for the subfamily. In addition, a minimum age constraint of 47.8 Ma was applied to the crown node of *Apeiba* based on the fossil *A. improvisa* MacGinitie (MacGinitie et al., 1974). Likewise, the crown node of *Luehea* was constrained to a minimum age of 41.2 Ma based on the fossil species *L. divaricatiformis* Fittip. et al., (Fittipaldi et al., 1989). An additional minimum age constraint of 47.8 Ma was applied to *Triumfetta* and *Heliocarpus* based on the fossil species *T. ovata* MacGinitie (MacGinitie, 1969). For fossil calibrations, only minimum age constraints were applied as fossils provide direct evidence of presence, but do not provide reliable information on the maximum age of the clade.

**Table 1.** Calibration constraints used in this study.

| Calibration/Fossil | Age (Ma) | Fossil Locality | Reference |
| --- | --- | --- | --- |
| Grewioideae crown | 64.41 | N/A | Hernández-Gutiérrez and Magallón, 2019 |
| *Apeiba improvisa*<br>MacGinitie | 47.8–<br>37.8 | USA | MacGinitie et al., 1974 |
| *Luehea divaricatiformis*<br>Fittip. et al. | 41.2–<br>33.9 | Brazil | Fittipaldi et al., 1989 |
| *Triumfetta ovata*<br>MacGinitie | 47.8–<br>37.8 | USA | MacGinitie, 1969 |

### 2.6. Diversification analyses

Diversification dynamics within *Grewia* were investigated using Bayesian Analysis of Macroevolutionary Mixtures (BAMM v.2.5.0) which reversible-jump Markov chain Monte Carlo (rjMCMC) to identify shifts in diversification regimes and estimate variation in speciation and extinction rates across time-calibrated phylogenies. Analyses were conducted on the dated *Grewia*-only phylogeny comprising 119 sampled species, which corresponds to a sampling fraction of 0.42 of currently recognized *Grewia* species (WFO, 2026). To account for incomplete and non-random taxon sampling, the expected number of species in each of the seven major *Grewia* clades we recovered was estimated using the morphological affinities and geographic distributions of unsampled taxa. When the placement of unsampled species could not be determined with confidence, species numbers were distributed among clades in proportion to their representation in the sampled phylogeny. The resulting clade-specific sampling fractions were 0.500, 0.436, 0.431, 0.432, 0.429, 0.432, and 0.433 for clades I–VII, respectively. A single diversification process was specified as the prior expectation (expectedNumberOfShifts = 1.0), and four Metropolis-coupled Markov (MCMC) chains were run for 50 million generations, sampling every 5,000 generations. Posterior samples were analysed using BAMMtools v.2.1.12, and the first 25% of samples were discarded as burn-in. Credible sets of diversification-rate configurations and posterior probabilities of rate-shift models were estimated using BAMMtools (Rabosky et al., 2014).

### 2.7. Geographical inference

The geographic distribution of each species included in the phylogeny was determined using floras: Flora of Tropical East Africa (Whitehouse, 2001), Flora Vitiensis (Seemann, 1873), regional treatments: Capuron (1964), Capuron and Mabberley (1999) Mabberley (1999), Chung (2005), Barrett (2019), and specimen data from major biodiversity databases, including the Global Biodiversity Information Facility (GBIF, 2026), iNaturalist (2026), Plants of the World Online (POWO, 2026), and World Flora Online (WFO, 2026). Additional distribution information was obtained from herbarium specimens examined by the authors in different herbaria (BR, MO, NY, P, US) and in digitalized collections (G, K).

Seven biogeographic regions were defined based on a combination of criteria including the present–day distribution of *Grewia*, the geological history and historical isolation of the major landmasses, and major barriers to plant dispersal (Cox et al., 2016; Torsvik and Cocks, 2017). These regions were A) Madagascar, B) continental Africa, C) western Indian Ocean islands, D) Arabia, E) Asia, F) Australia, and G) the Pacific (here represented by Fiji). Madagascar was distinguished from continental Africa because of its long geological isolation and distinctive biota (Goodman, 2022). Arabia was treated separately from continental Africa and Asia because of its distinct geological history and its transitional biogeographic position between the African and Eurasian biotas (Tejero-Cicuéndez et al., 2022; Ghazanfar, 2024). Asia and Australia were separated following the historical distinction between Sunda and Sahul (Hall, 2012, 2013; Crayn et al., 2015), and the Pacific was coded separately to distinguish dispersal into the oceanic Southwest Pacific (Gillespie and Clague, 2009; Holzmeyer et al., 2023). Comoros and Aldabra were grouped as western Indian Ocean islands to explicitly assess relatively recent over-water colonization events (Hume et al., 2018; Rougeau et al., 2025; Rusquet et al., 2025). Detailed species biogeographic assignments are provided in Supplementary File 1.

Ancestral ranges and historical biogeographic events were reconstructed using the R package BioGeoBEARS (Matzke, 2013, 2018) with the time-calibrated treePL tree inferred from only the *Grewia* sampling. We compared the DEC, DIVALIKE, and BAYAREALIKE models, each with and without the founder-event speciation parameter (J) (Matzke, 2014, 2022). The maximum number of areas per terminal was restricted to four. Model fit was evaluated using Akaike Information Criterion corrected for small sample size (AICc), and likelihood–ratio tests were used to assess the statistical support for alternative models. Biogeographical Stochastic Mapping (BSM) was subsequently performed under the best-fitting model to quantify the frequency and direction of inferred historical biogeographic events. A total of 1,000 stochastic maps were generated using the maximum-likelihood parameter estimates, and the resulting simulations were summarized to estimate the numbers of anagenetic dispersal, range contraction, founder-event speciation, vicariance, and within-area speciation events across the phylogeny (Matzke, 2016).

## 3. Results

### 3.1. Phylogenomic analysis

Target-capture sequencing yielded a highly complete dataset, with most samples recovering nearly all 353 targeted nuclear genes (mean = 349 (117–353) genes). After the exclusion of low-quality accessions, 171 samples were retained for phylogenetic analyses. After TrimAL exclusions, missing data were limited (7.86 % total gaps), indicating excellent locus recovery across the specimens sampled. After TrimAL and paralog removal, the final MSA was 142,580 bp long, spanning 342 loci.

Most nodes received strong SH-aLRT support (88.2% ≥ 80), with only a few shallow, recently diverged nodes having reduced support (11.8% < 80). Ultrafast bootstrap values were largely congruent (87.1% ≥ 95) with SH-aLRT though they generally provided stronger support at shallow nodes (i. e. UFBoot: 0% < 50). Site concordance factors averaged 52.6%, with 48.2% of internal branches showing sCF values ≥ 50% and only 4.1% showing values < 33.3%. Gene concordance factors were substantially lower, averaging 20.3%, with 11.8% of internal branches showing gCF values ≥ 50%, indicating considerable discordance among gene trees despite the high branch support in the concatenated phylogeny. The ASTRAL species tree was generally well supported, with 62.9% of internal branches receiving high local posterior probabilities (LPP ≥ 0.95), while only 5.9% showed low support (LPP < 0.50). Quartet support (q1) averaged 0.48, with 33.5% of internal branches receiving q1 values ≥ 0.50. All branches had q1 values above one-third, indicating that the topology selected by ASTRAL was consistently the most frequent of the three alternative quartet resolutions, despite substantial underlying gene-tree discordance.

We recovered two major clades in Grewioideae, *Apeibeae* and *Grewieae*, which had been resolved in prior studies (i.e., Brunken and Muellner, 2012, Dorr and Wurdack 2024, Hoorn et al., 2019, Colli-Silva et al., 2025). Within *Grewieae*, our results are congruent with those of Colli-Silva et al. (2025), and notably *Desplatsia* Bocq. is resolved as sister to *Grewia.* Note that *Eleutherostylis* Burret, which is resolved as the immediate sister group in Colli-Silva et al. (2025), is here treated as *Grewia* following Dorr and Wurdack (2024). *Grewia* was resolved as monophyletic with strong support (BS = 100%, Supplementary File 2). Our power to test the monophyly of *Grewia* is limited by our relatively sparse (generic level) Grewioideae sampling, but *Microcos* Burm. ex L. (sometimes considered congeneric with *Grewia*) is clearly well separated. The Malagasy species *G. erythroxyloides* Capuron occupies the earliest– diverging position within the genus. The remaining *Grewia* species are structured into seven clades (I–VII; Fig. 3). Clades I, IV, V, and VII received maximal support in both concatenated (SH-aLRT/UFBoot = 100/100) and coalescent analyses (LPP = 1). Clade V showed particularly high gene-tree concordance (gCF = 86.7%). In contrast, Clade III received comparatively low support in the concatenated analysis (SH-aLRT/UFBoot = 61/89; sCF = 32.3%) but was strongly supported by gene-tree and coalescent–based measures (gCF = 61.7%; LPP = 1; q1 = 0.86). Clade VI showed the opposite pattern, with maximal support in the concatenated analysis and high gene concordance (gCF = 75.9%), but lower support in the ASTRAL analysis (LPP = 0.81; q1 = 0.39).

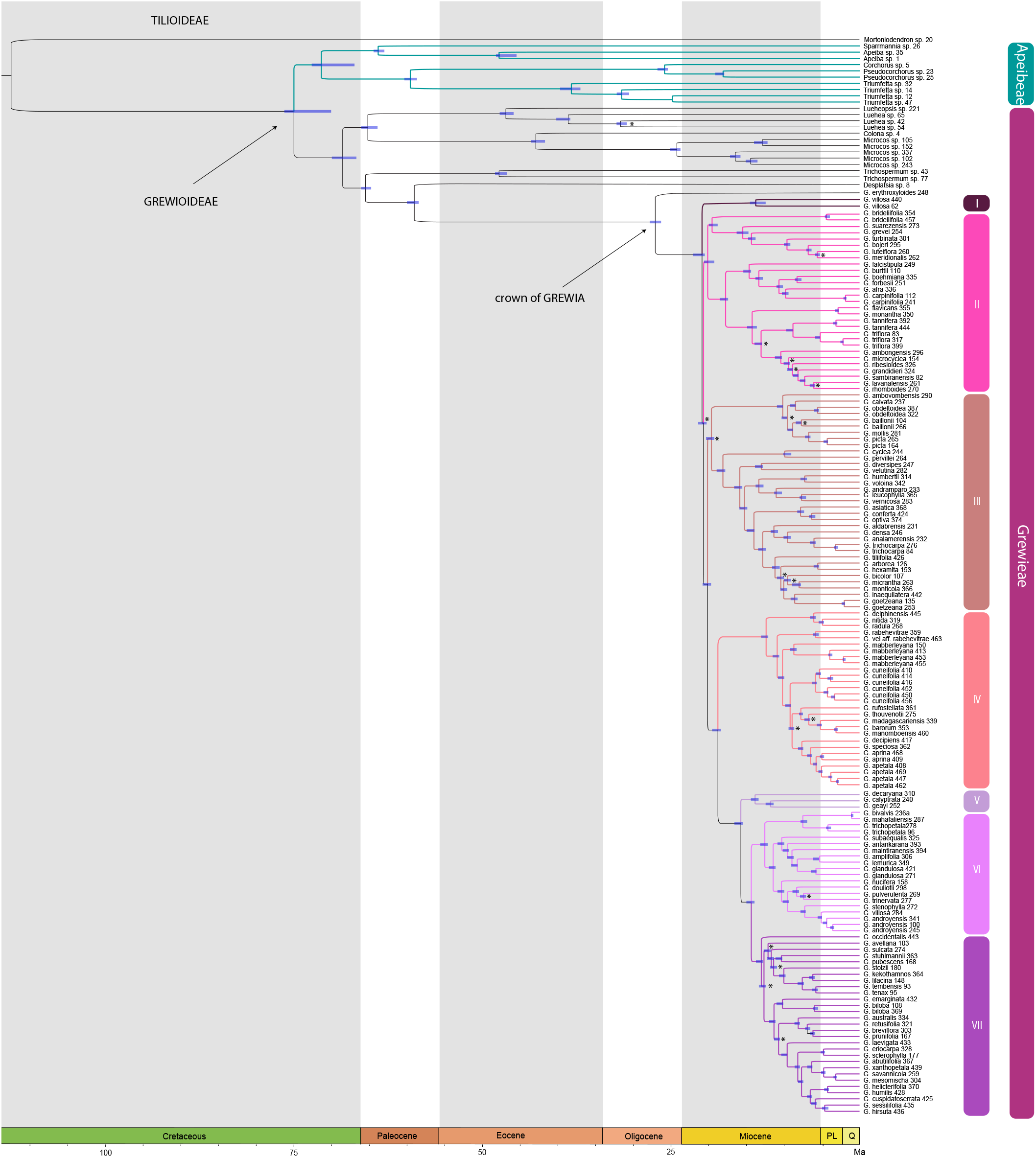

### 3.2. Divergence time estimation

Divergence time estimates obtained with treePL placed the crown age of subfamily Grewioideae in the Late Cretaceous at 74.81 Ma (67.69 – 75.00 Ma). Within the subfamily, the crown ages of Apeibeae and Grewieae were estimated at 71.36 Ma (65.89 – 71.56 Ma) and 68.55 Ma (64.31 – 69.62 Ma), respectively. A long stem branch of approximately 31.96 Ma separates the divergence of *Desplatsia* (59.04 Ma; 58.64 – 60.03 Ma) from the crown diversification of *Grewia* during the late Oligocene (27.08 Ma; 26.47 – 27.83 Ma). The seven major *Grewia* clades we identify here, subsequently originated within a relatively short interval from 20.89 Ma (Clade II), 19.64 Ma (Clade III) to 14.89 Ma (Clade VI), 13.85 Ma (Clade V), 13.76 Ma (Clade I), 13.02 Ma (Clade VII), and 12.39 Ma (Clade IV). Thus, all major lineages of *Grewia* were established within approximately 14 Ma following the origin of the crown group. Full age estimates and 95% confidence intervals of major clades are provided in Table 2 and Fig. 3.

**Table 2.** Age estimates (TreePL) with 95% CI, speciation, extinction, net diversification rates (BAMM) with 95% CI, and ancestral area reconstructions (R, BioGeoBEARS) for key nodes. Ages are Ma. Rates are species/Ma.

| Taxa | Age:<br>Mean<br>(95%<br>confidence<br>interval) | Speciation<br>rate:<br>Median<br>(95% CI) | Extinction<br>rate:<br>Median<br>(95% CI) | Net<br>diversification<br>rate: Median<br>(95% CI) | Ancestral area (probability) |
| --- | --- | --- | --- | --- | --- |
| Grewioideae | 74.81<br>(67.69 –<br>75.00) | – | – | – | – |
| Apeibeae | 71.36<br>(65.89 –<br>71.56) | – | – | – | – |
| Grewieae | 68.55<br>(64.31 –<br>69.62) | – | – | – | – |
| *Desplatsia* | 59.04<br>(58.64 –<br>60.03) | – | – | – | – |
| *Grewia* | 27.08<br>(26.47 –<br>27.83) | – | – | – | Madagascar (98.50%);<br>Africa (0.76%); Arabia<br>(0.75%) |
| Clade I | 13.76<br>(13.32 –<br>14.55) | 0.3426<br>(0.2806 –<br>0.4192) | 0.0236<br>(6.0e -04 –<br>0.0877) | 0.319<br>(0.2615 –<br>0.3807) | Africa (30.70%); Arabia<br>(26.19%); Africa + Arabia +<br>Asia (43.11%) |
| Clade II | 20.89<br>(20.41 –<br>21.56) | 0.2816<br>(0.2409 –<br>0.3326) | 0.022<br>(6.0e -04 –<br>0.0799) | 0.2596<br>(0.2221 –<br>0.2988) | Madagascar (99.95%) |
| Clade III | 19.64<br>(19.24 –<br>20.36) | 0.2683<br>(0.2299 –<br>0.3165) | 0.0218<br>(6.0e-04 –<br>0.0776) | 0.2466<br>(0.2097 –<br>0.2835) | Madagascar (99.99 %) |
| Clade IV | 12.39<br>(12.09 –<br>13.07) | 0.2335<br>(0.1962 –<br>0.2798) | 0.0222<br>(6.0e -04 –<br>0.0812) | 0.2113<br>(0.1717 –<br>0.2503) | Madagascar (99.98%) |
| Clade V | 13.85<br>(13.37 –<br>14.43) | 0.286<br>(0.2447 –<br>0.3376) | 0.0218<br>(6.0e -04 –<br>0.0800) | 0.2641<br>(0.2262 –<br>0.3037) | Madagascar (100%) |
| Clade VI | 14.89<br>(14.59 –<br>15.71) | 0.2369<br>(0.2006 –<br>0.2813) | 0.0216<br>(6.0e -04 –<br>0.0777) | 0.2154<br>(0.1762 –<br>0.2524) | Madagascar (100%) |
| Clade VII | 13.02<br>(12.43 –<br>13.38) | 0.2412<br>(0.2046 –<br>0.2857) | 0.0216<br>(6.0e -04 –<br>0.0776) | 0.2196<br>(0.1809 –<br>0.2563) | Africa (100%) |

### 3.3. Diversification analyses

BAMM analyses supported heterogeneous diversification dynamics within *Grewia*, with 89.0% of the posterior distribution favoring one or more diversification-rate shifts. A single-shift model dominated the posterior (70.0% of samples) and the credible set (74.7% posterior probability), with the most strongly supported shift representing an increase in diversification rate located early in the diversification of *Grewia*, on the branch subtending all sampled lineages except *G. erythroxyloides* (Fig. 4). Models without shifts accounted for only 11.0% of the posterior distribution, whereas models with two or more shifts represented 19.0%. Diversification rates also varied among the seven major clades (Table 2), with Clade I showing the highest estimated speciation (λ = 0.343) and net diversification rates (r = 0.319), and Clade IV the lowest (λ = 0.234; r = 0.211). Extinction rates showed comparatively little variation among clades (μ = 0.022 – 0.024), indicating that differences in net diversification were primarily associated with variation in speciation rates.

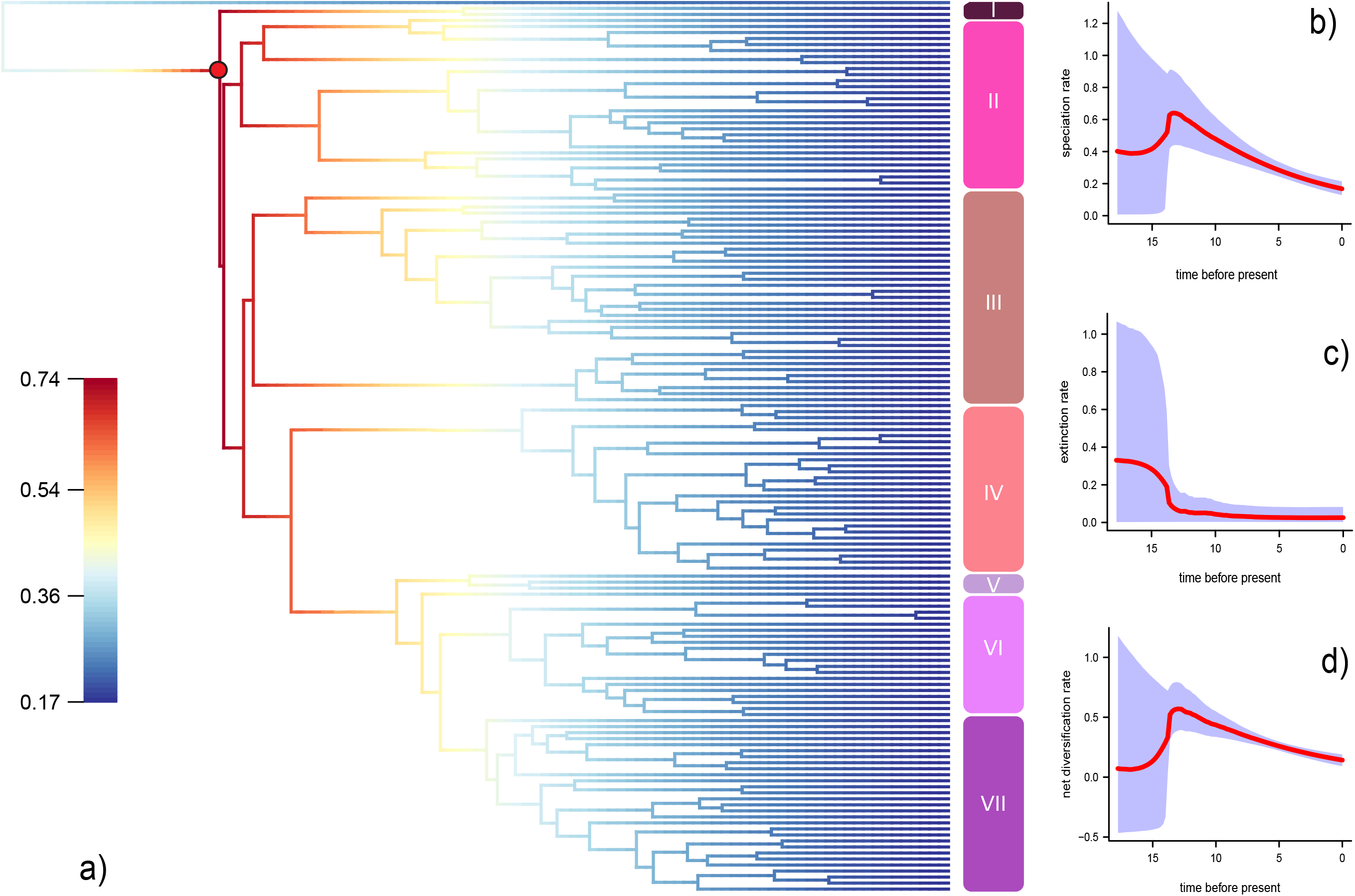

### 3.4. Geographic distribution

Model comparison strongly supported BAYAREALIKE+J as the best-fitting model, with the lowest AICc value and the highest Akaike weight (Table 3). Geographic structuring was evident across the seven major clades of *Grewia*. Malagasy species were distributed across six of the seven clades and occupied early diverging positions within five of those lineages. Clades IV and V are composed of species restricted to Madagascar. Asian species are restricted to Clades III and VII, whereas Arabian species occurred in clades that also contained African taxa (Clades I, II, III, VI and VII). Species from the Comoros and Aldabra were only recovered in clades containing Malagasy species (Clades II, III, and VI). Australian species, together with the accession from the Pacific, were confined to Clade VII. In Clade III, lineages dispersed from Madagascar to the Arabian Peninsula (ca. 13.0 Ma) and subsequently to Asia (ca. 14.1 Ma), where further diversification occurred. Our reconstruction further suggests a secondary dispersal from Asia back to continental Africa (ca. 11.2 Ma).

**Table 3.** Comparison of the biogeographic models implemented in BioGeoBEARS.

| Model | LnL | k | ΔAICc | AICc | AICw | d | e | j |
| --- | --- | --- | --- | --- | --- | --- | --- | --- |
| DEC | -212.354 | 2 | 75.734 | 428.791 | 3.58e - 17 | 0.002 | 1.00e - 12 | — |
| DIVALIKE | -212.758 | 2 | 76.544 | 429.600 | 2.39e - 17 | 0.002 | 1.00e - 12 | — |
| BAYAREALIKE | - | 2 | 129.80 | 482.863 | 6.50e - | 0.002 | 0.033 | — |
|  | 239.3<br>89 |  | 7 |  | 29 |  |  |  |
| DEC J | -<br>197.0<br>88 | 3 | 47.287 | 400.343 | 5.39e -<br>11 | 1.00e -<br>12 | 1.00e -<br>12 | 0.011 |
| DIVALIKE J | -<br>201.5<br>11 | 3 | 56.134 | 409.189 | 6.47e -<br>13 | 1.00e -<br>12 | 1.00e -<br>12 | 0.010 |
| **BAYAREALIKE J** | **-**<br>**173.4**<br>**44** | **3** | **0** | **353.056** | **1.00** | **1.00e -**<br>**12** | **1.00e -**<br>**12** | **0.014** |

Shown are the log-likelihood (LnL), number of free parameters (k), estimated dispersal (d), extinction (e) and founder-event speciation (j) parameters, corrected Akaike Information Criterion (AICc), $\Delta$ AICc, and Akaike weights (AICw). The best-supported model is highlighted in bold.

### 3.5. Biogeographic stochastic mapping

Biogeographic stochastic mapping indicated that the historical biogeography of *Grewia* was overwhelmingly shaped by dispersal, with founder-event speciation (21.39 ± 0.67) and range expansion (12.29 ± 0.59), whereas vicariance and other cladogenetic processes were essentially absent. The highest number of dispersal events was inferred from Madagascar to continental Africa (13.38 ± 1.24), identifying Madagascar as the principal source of westward colonization. Africa subsequently emerged as a secondary hub, with frequent dispersal toward Arabia (5.31 ± 0.99) and Asia (4.30 ± 1.00), whereas Asia represented the main source of colonization for Australia (4.62 ± 0.48). Additional exchanges between Madagascar and the western Indian Ocean islands, together with recurrent movements between Africa and Madagascar, further highlight the dynamic connectivity of the western Indian Ocean (Table 4, Fig. 5).

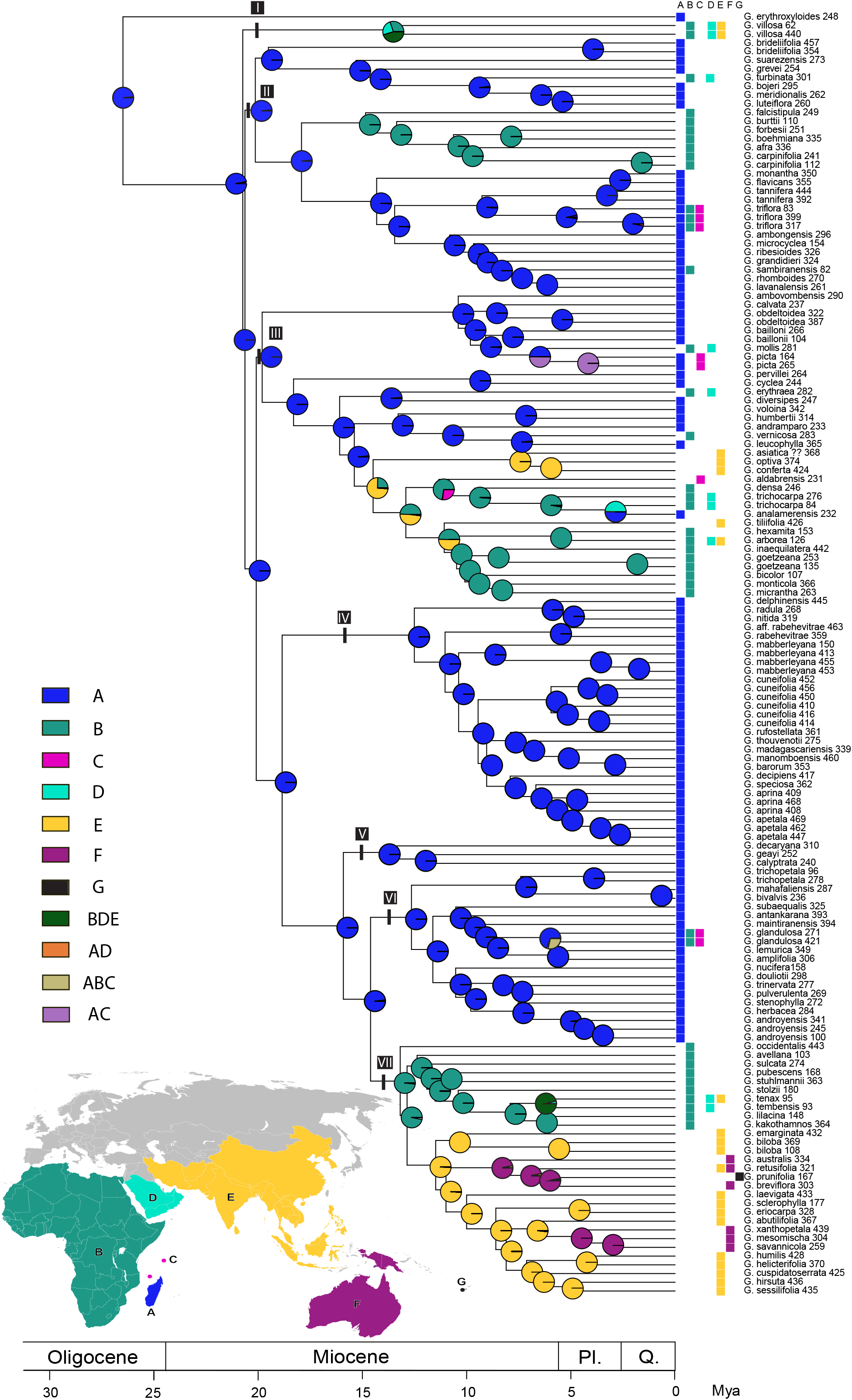

**Table 4.**
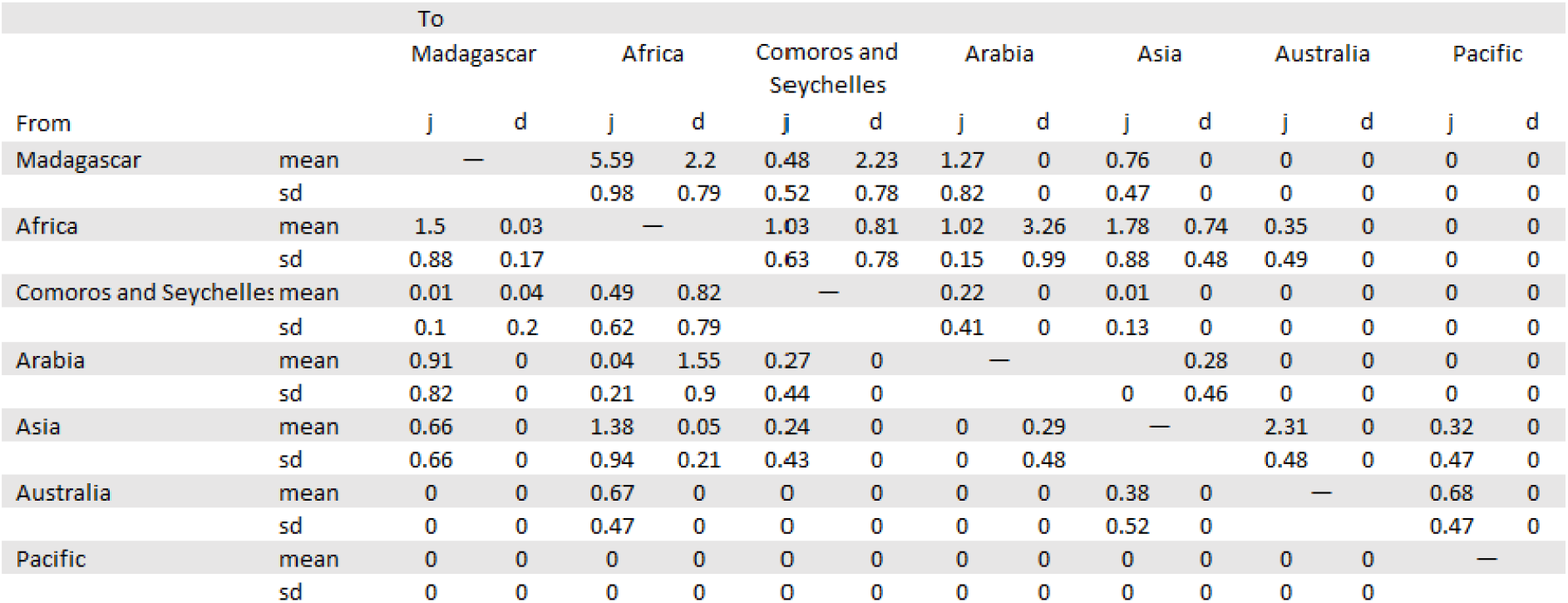
Summary of dispersal events. Events were calculated and averaged over 1000 BSMs in BioGeoBEARS. d: anagenetic dispersal; j: founder-event speciation.

## 4. Discussion

### 4.1. First genomic phylogeny of Grewia

We present the first phylogenomic study of *Grewia* and further demonstrate the utility of Angiosperms353 for resolving relationships within complex, species-rich groups. The previous most comprehensive *Grewia* phylogeny sampled 61 species but used only a single locus, ITS (Dorr and Wurdack, 2024). Although only 35 *Grewia* spp. are shared between the two studies, there is general congruence among the major clades (in grouping and ordering) except our Clades II and III are reversed order in that study. *Desplatsia*, with four species from tropical West and Central Africa, is the sister group to species-rich *Grewia,* demonstrating unequal diversification dynamics*. Desplatsia* resembles *Grewia* in general morphology but notably differs in large (to 20 – 25 cm) elephant dispersed fruit (Wellsow et al., 2019). *Microcos* has been united with *Grewia* in the past (Wight and Walker-Arnott 1834; Drummond, 1915), but it is morphologically distinct with differences in leaf margins, inflorescence structure, bracteoles, and androgynophores and our work supports maintaining both as distinct genera. *Grewia erythroxyloides* Capuron, from southern Madagascar is the unexpected earliest-diverging lineage in the genus and is morphologically relatively specialized as a shrub with lateral short shoots of small obovate leaves in fasciculate clusters (i.e., brachyblasts), and yellow flowers (Fig 1. e). Its exact resolution may be affected by incomplete taxon sampling but not by obvious data quality issues (348 genes recovered with 0.0098 % of gap proportion for this tip)

The major clades recovered here broadly correspond to some previously proposed sections of *Grewia* (Capuron, 1964; Capuron and Mabberley 1999; Mabberley, 1999). Clade II includes Malagasy representatives of subg. *Burretia* (Hochr.) Capuron and subg. *Vincentia* (Benth.) Capuron, together with African and Arabian species, whereas Clade III comprises Asian, African, Arabian, and Malagasy species, including a few representatives of sect. *Axillares* Burret. Although broader sampling and explicit analyses of morphological character evolution will be needed to assess these patterns, our phylogenetic framework provides a basis for testing existing classifications and developing a much-needed new robust infrageneric classification of *Grewia*.

### 4.2. Madagascar is the center of Grewia diversity

The ca. 32 Ma stem branch separating the origin of *Grewia* from the crown diversification suggests that early-diverging stem lineages may have gone extinct before the radiation of extant species (Quental and Marshall, 2010; Budd and Mann, 2020). The ancestral area reconstruction strongly supports Madagascar as the ancestral area of crown *Grewia* (98.5%; Table 2; Fig. 5). However, *Desplatsia*, the sister lineage to *Grewia*, is restricted to Africa, raising the possibility that the common ancestor of the two lineages occurred in Africa, followed by dispersal to Madagascar along the stem lineage of *Grewia* and subsequent diversification on the island.

The crown diversification of *Grewia* is estimated to have occurred during the late Oligocene to early Miocene around 27.08 Ma (26.47 – 27.83 Ma), well after the separation of Madagascar from Africa beginning around 170 Ma (Reeves, 2014), and its subsequent separation from India around 88 Ma (Gibbons et al., 2013; Reeves, 2014). Thus the origin of the genus cannot be explained by Gondwanan vicariance and instead reflects successful overseas dispersal followed by in situ diversification, a biogeographic scenario that has also been inferred for diverse Malagasy plant and animal lineages (Warren et al., 2010; Bacon et al., 2016; Masters et al., 2021; Génin et al., 2022; Ali and Hedges, 2023; Skema et al., 2023).

The diversification-rate shift inferred in *Grewia*, dated to ca. 25 Ma, occurred while the early diversification of the genus was still centered in Madagascar, before its first expansions into Africa and Arabia (Clade I). The timing of this marked increase in diversification rate in *Grewia* falls in the Oligocene Warming, an interval characterized by increasing sea-surface temperatures (O’Brien et al., 2020; Guitián et al., 2021) but also substantial fluctuations in Antarctic ice volume (Pekar et al., 2006; Brzelinski et al., 2023). It also falls within the broader Cenozoic renewal of angiosperm diversification (Dimitrov et al., 2023). This early radiation was followed by extensive exchange between Madagascar, continental Africa and the western Indian Ocean islands. Water-circulation models suggest that dispersal across the Mozambique Channel was particularly favorable during the Paleogene and early Miocene, before a reorganization of ocean currents around 20–15 Ma progressively reduced the likelihood of eastward dispersal (Ali and Huber, 2010; Crottini et al., 2012).

Our ancestral reconstructions reveal a marked asymmetry in dispersal direction across the Mozambique Channel. Biogeographical stochastic mapping recovered Madagascar-to-Africa as the dominant dispersal route, with an average of 5.59 ± 0.98 founder-event dispersals and 0.03 ± 0.17 anagenetic range expansions, compared with only 1.50 ± 0.88 founder events and 0.03 ± 0.17 anagenetic dispersals from Africa to Madagascar. With our current sampling we identify only one species (*Grewia analamerensis* Capuron, from Clade III) known from Madagascar with their closest living relatives in mainland Africa/Arabia. Within Clade II, we reconstruct two westward dispersal events from Madagascar to continental Africa, inferred around 14.2 Ma and 5.2 Ma. Likewise, the reconstruction of an African ancestral area for the crown of Clade VII (ca. 13 Ma) is ultimately derived from an earlier colonization of the continent from Madagascar, followed by subsequent diversification within Africa. These temporal patterns closely match broader biogeographic trends documented in Malagasy mammals. Crottini et al. (2012) inferred a predominance of dispersal events out of Madagascar into Africa since ca. 30 Ma, with a marked increase during the Miocene (ca. 10– 20 Ma), broadly coinciding with our results.

The dispersal of *Grewia* to western Indian Ocean islands shows a different pattern. Founder-event speciation associated with island colonization from Madagascar to Comoros and Aldabra was uncommon (0.48 ± 0.52 events), whereas anagenetic dispersal involving these islands was more frequent (2.71 ± 0.78 events). The recent inferred colonization of the Comoros that we recover is consistent with the earliest well-documented subaerial volcanism in the region around 2.4 Ma (Rusquet et al., 2025). Within Clade II, the ancestral distribution of the three samples of *G. triflora* (Bojer) Walp. was reconstructed as Madagascar with very high support (99.99%), suggesting that the species’ present occurrence in the Comoros and continental Africa likely resulted from recent dispersal from Madagascar. Similarly, samples of *G. glandulosa* Vahl (Clade VI) retained a predominantly Malagasy ancestral area (76.1%) at 5.78 Ma despite its current occurrence in continental Africa and Aldabra. Other species with current distributions in Madagascar, Aldabra, and continental Africa (Clade III) likewise retained a substantial Malagasy ancestral component. Although our reconstructions do not indicate that the Comoros and Aldabra acted as stepping stones for the inferred dispersal events from Madagascar to Africa, geological reconstructions suggest that the Mozambique Channel was far more topographically complex in the past than today. Volcanic ridges, episodes of uplift, and now-submerged topographic highs may have intermittently reduced dispersal distances across the channel during the Cenozoic (Masters et al., 2021; Aslanian et al., 2023).

Our work adds *Grewia* to a growing list of plant lineages with documented westward dispersal from Madagascar to Africa and/or into the western Indian Ocean islands (Krüger et al., 2012; Strijk et al., 2012; Skema et al., 2023; Wan et al., 2024). Notably, no species of *Grewia* have been recorded in the Mascarene Islands. This absence is surprising given recent documentation of colonization events from Madagascar to the Mascarenes in other plant groups, including Malvaceae (e.g. Le Péchon et al., 2015; Strijk et al., 2012). Whether this reflects dispersal limitation, ecological filtering after arrival, or subsequent extinction remains unclear.

### 4.3. An Afro-Arabian corridor

After *Grewia erythroxyloides*, the earliest-diverging lineage in *Grewia* (Clade I) was reconstructed as having an Afro-Arabian ancestral distribution at 13.76 Ma, with ancestral state probabilities of 43.11% for Africa + Arabia + Asia, 30.70% for Africa alone, and 26.19% for Arabia alone. Beyond this early divergence, Arabian lineages were consistently reconstructed as originating from adjacent regions rather than diversifying in situ. Although this pattern should be interpreted cautiously given the limited number of Arabian species included in our sampling, the BSM analyses recovered Africa as the principal source of dispersal into Arabia (3.26 ± 0.99 anagenetic and 1.02 ± 0.15 founder-event dispersals), whereas movements in the opposite direction were much less frequent (1.55 ± 0.90 anagenetic and 0.04 ± 0.21 founder-event dispersals). Similarly, dispersals from Africa to Asia (0.74 ± 0.48 anagenetic and 1.78 ± 0.88 founder-event events) exceeded the reverse direction.

A variety of angiosperm lineages have reconstructed dispersal events between Asia and Africa that coincide with major tectonic and paleogeographic changes during the Early to Middle Miocene (e.g., Zhou et al., 2012; Yu et al., 2014; Yang et al., 2016). The progressive closure of the Tethyan Seaway and the collision of the Afro-Arabian and Eurasian plates established terrestrial connections between Afro-Arabia and Eurasia, including the temporary Gomphotherium Landbridge at ca. 18–16 Ma (Rögl, 1999; Portik and Papenfuss, 2015), followed by a more permanent land connection from ca. 15 Ma onwards (Portik and Papenfuss, 2015). These connections facilitated extensive biotic exchanges between Africa and Eurasia (Harzhauser et al., 2007; Sen, 2013; Portik and Papenfuss, 2015) and correspond with the dispersal events and estimated time frames we recover in *Grewia*.

The Middle Miocene Climatic Optimum represented the warmest interval of the Miocene, with generally warmer and more humid conditions that promoted the expansion of vegetation (Steinthorsdottir et al., 2021). Fossil and paleoenvironmental evidence further suggest that Arabia supported open woodland, bushland, and locally mesic habitats during much of the Miocene, in strong contrast to the hyper-arid environment that defines much of the region today (Markowska et al., 2025). This paleoenvironmental framework lends support to the dispersal events we infer in *Grewia* between Africa and Arabia. Furthermore, dispersal from Africa into Asia during the Miocene has been inferred in several other plant lineages, e.g. in *Uvaria* L. (Annonaceae), with dispersal through Arabia during the Miocene climatic optimum (Zhou et al., 2011), and in *Indigofera* L. (Fabaceae), with multiple dispersal events from Africa into Asia (Du Preez et al., 2025).

### 4.4. From Sunda to Sahul and to the Southeast Pacific

Clade VII was reconstructed as originating in continental Africa. It subsequently dispersed to tropical Asia ca. 13.0 Ma and to Australia within the last 10 Ma. This timing coincides with the progressive northward convergence of the Australian Plate with Southeast Asia ca. 12 Ma (Hall, 2009, 2012). The resulting tectonic reorganization progressively reshaped Wallacea and reduced marine distances between the Sunda and Sahul shelves, facilitating biotic exchange between Asia and Australia (Hall, 2009, 2012; Baldwin et al., 2012). The island chain facilitated floristic exchange in both directions but was strongly asymmetric, with dispersal from Sunda to Sahul (Richardson et al., 2012; Crayn et al., 2015; Peng et al., 2021; Holzmeyer et al., 2023) substantially more frequent than in the opposite direction (Wu et al., 2018). In *Grewia*, we recovered 2.31 (± 0.48) founder event dispersals from Asia to Australia and 0.38 (± 0.52) founder events in the reverse direction.

The single Pacific representative of *Grewia* in our study, *G. prunifolia* A.Gray from Fiji, was reconstructed as deriving from a founder event, most frequently from an Australian ancestor (j=0.68 (± 0.47)), although an Asian origin was also recovered (j=0.32 (± 0.47)). Both scenarios are compatible with dispersal through New Guinea and the Solomon Islands to Fiji, a pathway inferred for several plant lineages (Johnson et al., 2017; Swenson et al., 2019; Holzmeyer et al., 2023). However, broader sampling of Clade VII, particularly from New Guinea and Australia (ca. 20 spp. occuring), will be necessary to clarify the origin and dispersal route of the Pacific lineage.

### 4.5. Attractive fruits facilitating dispersal

The fruit of *Grewia* are generally small (0.3 – 4.5 cm), entire or lobed drupes (Fig. 1). A fleshy-fibrous pericarp encases the seeds that are in turn well protected within pyrenes and lignified seed coats (Corner, 1976). The sweet fruits of multiple *Grewia* species (e.g., *G. asiatica* L., *G. flavescens* Juss., *G. tenax* (Forrsk.) Fiori) are regularly foraged and/or cultivated for human consumption and are rich in minerals and antioxidants (Gebauer et al., 2013; Islary et al., 2016). These fruit characteristics are consistent with vertebrate-mediated endozoochory (Brunken and Muellner, 2012) and frugivory has been reported for at least ten *Grewia* species across Africa, Asia, and Australia (Fricke and Svenning, 2020). Documented consumers of the genus *Grewia* include primates (Wrangham and Waterman, 1983; Spehn and Ganzhorn, 2000), elephants (Baskerville et al., 2011; Baskaran and Desai, 2013), fruit bats (Cumming and Bernard, 1997; Fricke and Svenning, 2020), canids (Favaretto et al., 2024), tortoises and lizards (Fricke and Svenning, 2020), as well as numerous birds, including hornbills (Kannan and James, 1999), parrots (Symes and Perrin, 2003), mousebirds (Rowan, 1967), bulbuls, mynas, and other frugivorous species (Fricke and Svenning, 2020). In Madagascar, *Grewia* fruits are known to be a major diet component for lemurs and studies have further explored dispersal, seed germination rates (with seeds recovered from feces) and even ‘gardening’ of *Grewia* by lemurs (Spehn and Ganzhorn, 2000; Sato, 2012; Génin and Rambeloarivony, 2018). Although these contemporary observations cannot identify the vectors responsible for the historical dispersal events reconstructed here, they indicate that *Grewia* fruits are currently exploited by a remarkably broad diversity of vertebrates and that their seeds remain viable after gut passage, providing a biologically plausible mechanism for both regional seed dispersal and occasional long-distance dispersal.

## 5. Conclusions

The assembly and diversification of the Paleotropical flora has been profoundly shaped by plate tectonic processes, climatic fluctuations (especially changes in aridity), and the progressive reconfiguration of tropical land connections. Much of this dynamic history helps explain our biogeographic and temporal reconstructions in *Grewia*. By integrating a densely sampled phylogenomic framework with divergence-time estimation, diversification analyses, and probabilistic biogeographic reconstructions, our study reveals a complex evolutionary history of *Grewia* across the Paleotropics. These results strengthen our understanding that Madagascar can act as a source for lineages and not merely as a place of in situ diversification. We recover a Malagasy origin for the crown of the genus *Grewia* at ca. 27 Ma, followed by a single shift toward increased diversification rates at ca. 25 Ma, when the lineages were still mainly restricted to Madagascar. This early diversification was followed by repeated dispersal events that successively connected Madagascar with continental Africa, the western Indian Ocean islands, Arabia, tropical Asia, Australia and the southwest Pacific.

*Grewia* fruits are adapted for animal consumption and provide a plausible mechanism to explain its large distribution and much of the overland and over sea dispersal we reconstruct for the genus. Our findings in *Grewia* further highlight the importance of geologic and climatic fluctuations in the Miocene, especially in and around the Mozambique Channel, Afro-Arabia, and throughout the Sunda, Wallacea, and Sahul regions to the development and distribution of the Paleotropical flora.

## CRediT authorship contribution statement

Lucile Jourdain-Fievet: Conceptualization, Data curation, Formal analysis, Funding acquisition, Investigation, Methodology, Project administration, Resources, Software, Validation, Visualization, Writing – original draft, Writing – review and editing.

Margaret M. Hanes: Conceptualization, Methodology, Resources, Supervision, Validation, Writing – review and editing.

Nisa Karimi: Conceptualization, Formal analysis, Methodology, Resources, Supervision, Writing – review and editing.

Laurence J. Dorr: Conceptualization, Funding acquisition, Investigation, Methodology, Resources, Supervision, Validation, Writing – review and editing.

Kenneth J. Wurdack: Conceptualization, Funding acquisition, Investigation, Methodology, Resources, Supervision, Validation, Writing – review and editing.

## Data availability

Raw sequence reads in FASTQ format will be deposited in the NCBI Sequence Read Archive (accession XXX) after review.

## Acknowledgements

We thank several herbaria (G, MO, NY, and US), for providing leaf samples; P and BR for granting access to their collections; Gabriel Johnson for laboratory support; Laurence Ramon for advice and identification of some specimens; Rose A. Gulledge for Fig. 1; and Rebeca Hernández-Gutiérrez for sharing fossil data. The laboratory work and data analyses were conducted in and with the support of the Laboratories of Analytical Biology of the National Museum of Natural History, Smithsonian Institution.

We also thank Riana Fourie, Graham Grieve, Neha Jaiswal, Dewald du Plessis, Marco Schmidt, Russell Cumming, Mo Haibo, Nuwan Jayawardana, Alexander Ivanov, and Heath Milne for kindly allowing us to use their photographs available through iNaturalist in Fig. 1.

## Appendix A. Supplementary data

Supplementary File 1. Table of vouchers origins and geographic coding.

Supplementary File 2. Phylogenomic supports (SH-aLRT; UFBoot; gCF; sCF; LPP; quartets).

